# Comparing the efficiency of six clearing methods in developing seeds of *Arabidopsis thaliana*

**DOI:** 10.1101/2022.09.12.507557

**Authors:** Venkata Pardha Saradhi Attuluri, Juan Francisco Sánchez López, Lukáš Maier, Kamil Paruch, Hélène S. Robert

## Abstract

Tissue clearing methods eliminate the need for sectioning, thereby helping better understand the 3D organization of tissues and organs. In the past fifteen years, clearing methods have been developed to preserve endogenous fluorescent protein tags. Some of these methods (ClearSee, TDE, PEA-Clarity, etc.) were adapted to clear various plant species, with the focus on roots, leaves, shoot apical meristems, and floral parts. However, these methods have not been used in developing seeds beyond the early globular stage. Tissue clearing is problematic in post-globular seeds due to various apoplastic barriers and secondary metabolites. In this study, we compared six methods for their efficiency in clearing *Arabidopsis thaliana* seeds at post-globular embryonic stages. Three methods (TDE, ClearSee, and ClearSee alpha) have been already reported in plants whereas the others (fsDISCO, FAST9, and CHAPS clear) are used in this context for the first time. These methods were assessed for seed morphological changes, clearing capacity, removal of tannins, and spectral properties. We tested each method in seeds from globular to mature stages. The pros and cons of each method are listed herein. ClearSee alpha appears to be the method of choice as it preserves seed morphology and prevents tannin oxidation. However, FAST9 with 60% iohexol as a mounting medium is faster, clears better, and appears suitable for embryonic shape imaging. Our results may guide plant researchers to choose a suitable method for imaging fluorescent protein-labeled embryos in intact Arabidopsis seeds.

**Key message:** ClearSee alpha and FAST9 were optimized for imaging Arabidopsis seeds up to the torpedo stages. The methods preserve the fluorescence of reporter proteins and seed shape, allowing phenotyping embryos in intact seeds.

## Introduction

Seed development is an essential process in the life cycle of angiosperms. It is initiated by double fertilization, which leads to the development of the embryo and endosperm. While the endosperm acts as a nutritional source for the developing embryo, the embryo develops into a miniature seedling embedded in the seed, ready to germinate when conditions are optimal. The seed coat comprises several integument layers protecting the embryo from various biotic and abiotic factors (Supplementary Fig. 1) (Matilla 2019; Doll and Ingram 2022; Verma et al. 2022). Plant developmental researchers use embryos for the study of body pattern formation. However, the seed coat often needs to be removed to visualize expression patterns of embryonically expressed genes. Alternatively, the seeds may be sliced into thin sections for whole seed imaging. Both methods remove the embryo from its spatial context, which might complicate the understanding of diverse developmental aspects.

Methods with deeper imaging capability of the plant tissues without sectioning organs are invaluable in developmental biology. Most plant tissues have a limitation of about 50 μm imaging depth due to light scattering imposed by the difference in the refractive index of cellular structures (Hériché et al. 2022). Tissue clearing methods help minimize these differences, thus enabling deeper imaging of the plant tissues. However, each clearing method has a specific protocol for clearing the tissue. Although the first tissue clearing methods are older than one hundred years (Hoyer 1882), clearing methods developed in the past two decades are compatible with genetically expressed fluorescent proteins (FPs) and different dyes and stains (Hériché et al. 2022). Clearing thick tissues and organs while preserving the fluorescence of dyes and FPs opens up new possibilities for studying the development of various organs with genetically expressed fluorescence markers.

Recently, some clearing methods developed for animals were successfully applied in plant tissues (ClearSee, ClearSee alpha, thiodiethanol [TDE], PEA-CLARITY, and a urea-based method) with or without modifications. Warner et al (2014) used a urea-based clearing medium on tissues and organs from various plant species, including root nodules from pea. TDE clearing was successfully applied in plant tissue (Musielak et al. 2016) using a mixture of TDE and water, which worked in *Arabidopsis* pistil and young seeds. ClearSee uses bile salts, xylitol for clearing and decoloration of tissue, and urea for refractive index matching (Kurihara et al. 2015). The method was further improved (and named ClearSee alpha) by adding antioxidants in the clearing solution to prevent the browning of tissues with high anthocyanin content (Kurihara et al. 2021). On the other hand, PEA-CLARITY uses acrylamide for tissue transformation and SDS for clearing. Cell-wall digestion of PEA-CLARITY-transformed tissue is compatible with immunolocalization (Palmer et al. 2015).

In this study, we investigated three methods that were not previously reported for plant tissues (fsDISCO, CHAPS clear, and FAST9). The fsDISCO clearing method is a variation of 3DISCO, an organic solvent-based method known for its high clearing capabilities in a short time (Ertürk et al. 2012). 3DISCO is a simple two-step process. The first step involves organ dehydration using graded tetrahydrofuran (THF) and the second step involves matching the refractive index using dibenzyl ether (DBE). However, the original 3DISCO quenched the fluorescence of FPs and was limited to samples with a high fluorescent signal. The recent modifications to the 3DISCO protocol, called fDISCO (fluorescent compatible) and sDISCO (stabilized DISCO), eliminated the drawbacks of the 3DISCO by retaining the fluorescence of FP and increasing photostability, respectively (Qi et al. 2019). fDISCO retains fluorescence of FPs by allowing the tissue dehydration at low temperature with pH-adjusted solvents. These two modifications are compatible with all FPs tested and increased fluorescence compared to 3DISCO. sDISCO makes FPs more photostable by using free radical scavengers (Hahn et al. 2019). We have combined these methods to achieve higher fluorescence and more photostability, calling this method fsDISCO.

FAST9 (**F**ree of **a**crylamide **s**odium dodecyl sulfate (SDS)-based **t**issue clearing with pH**9**) is a simplified version of a method called SHIELD. Like PEA-CLARITY, SHIELD uses a gel matrix to hold biomolecules and SDS to remove lipids (Park et al. 2019). Later, the refractive index of the cleared tissue is matched with EasyIndex (LifeCanvas technologies), a proprietary high refractive index matching formulation (https://lifecanvastech.com/products/easyindex/). Although the workflow resembles CLARITY, there is a significant difference, namely using an epoxy resin to form a hydrogel matrix instead of an acrylamide one. The resin successfully protects FPs from temperature, pH, and chemical exposure (Park et al. 2019). Our initial trials with SHIELD showed FP protection in the plant tissue; however, we observed oxidation of the tannins, which makes this method unsuitable for seed clearing.

Some tissues are more challenging to clear due to their compact extracellular molecular mesh. Detergents like SDS (present as micelles) do not perform well as they cannot get freely in and out of the tissue. To overcome this challenge, SHANEL, another clearing method, uses CHAPS as a detergent to improve tissue permeability (Zhao et al. 2020). CHAPS makes smaller micelles than SDS, making it suitable for clearing the tissue with the abovementioned properties. We reasoned that CHAPS could be used for clearing plant tissue (CHAPS Clear).

This article focuses on the strengths and weaknesses of each clearing method by comparing their efficiency in removing tannins, tissue clearing, morphological preservation, and spectral properties after clearing. We use *Arabidopsis* developing seeds for the testing because only few methods are available for this tissue. We believe our findings may help plant researchers choose appropriate methods and help those who want to develop new methods.

## Results

### The tannin removal efficiency varies depending on the method

Proanthocyanidins (PAs), also called condensed tannins, are colorless flavonoid polymers in the inner integument of Arabidopsis (Supplementary Fig. 1). They confer the brown seed color, after oxidation, during the mature stages of the seed development. During seed development, PAs accumulate (mainly in the inner integument 1 [ii1] layer) from the globular to heart stages and continue through the green cotyledon stage (Debeaujon et al. 2003). The accumulation is non-uniform and starts from the micropylar pole to end at the chalaza (Debeaujon et al. 2003). Oxidized PAs in the seed coat serve as light filters, attenuators modifying the spectra of the incident light to protect the embryo from radiation damage.

However, this attenuating property is not desirable for imaging purposes. We used TDE and ClearSee protocols to clear seeds containing globular to heart stage embryos. Our initial studies found that the oxidized tannins effectively block the 405nm emission wavelength preventing clean imaging of the embryo in Renaissance SR2200-stained seeds (Supplementary Fig. 2). Thus, we reasoned that removal of PAs would render a better clearing.

We stained the cleared seeds with vanillin (Xuan et al. 2014) to assess the ability of the tested clearing methods to remove tannins. Since the tannin accumulation is not uniform during the early embryonic stages, we choose to stain the seeds with vanillin five days after pollination (dap, hand-pollinated; heart stage) to test a developmental stage known to accumulate PAs in all seed coat regions. The results are presented in Fig. 1.

**Fig. 1.**
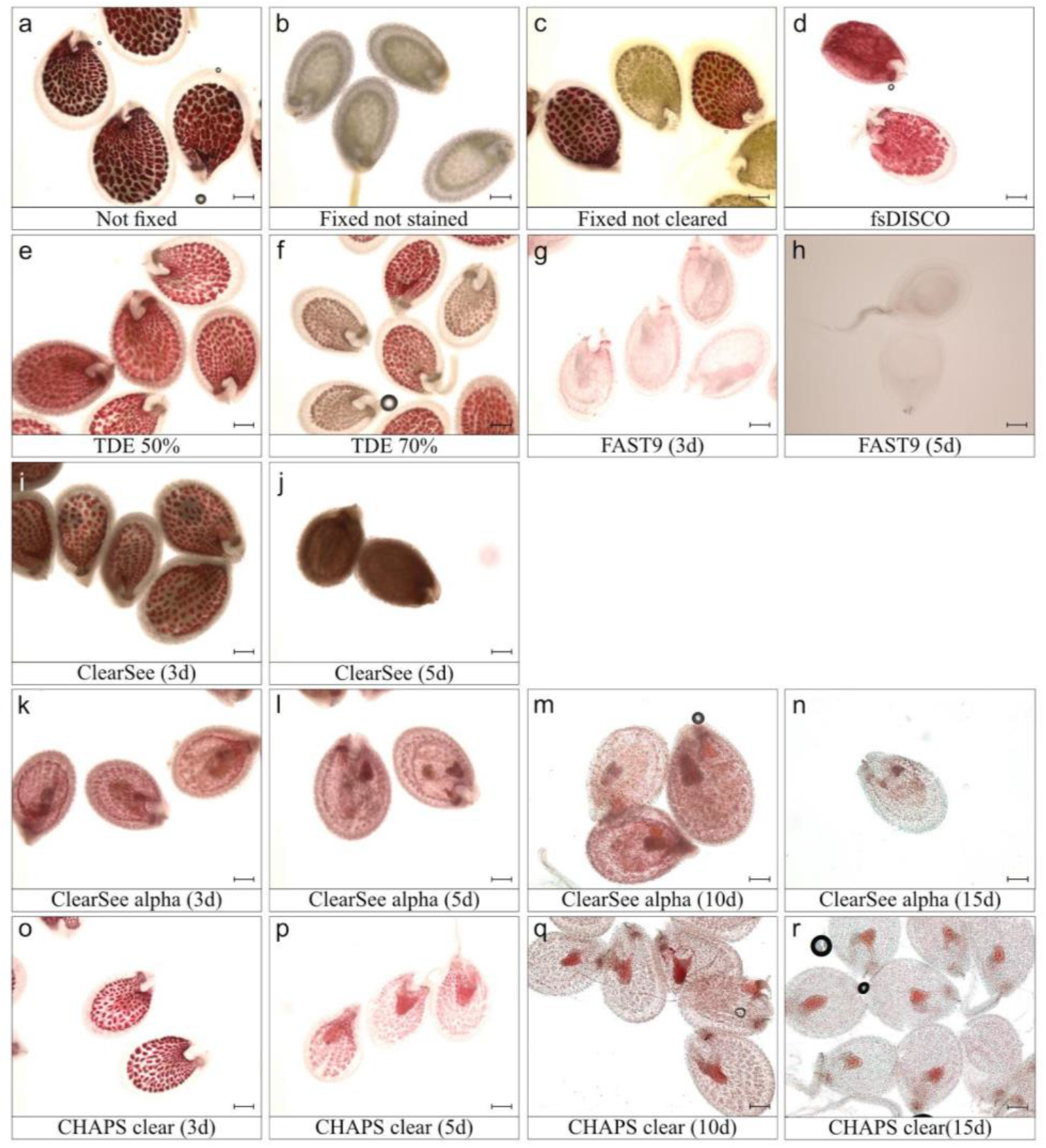
Vanillin staining of Arabidopsis seeds cleared with different methods. Vanillin staining tests the presence of tannins after clearing. (a-c) Control seeds: not fixed (a), fixed and not stained with vanillin (b), fixed and stained (c). (d-r) Seeds cleared with the different clearing methods tested: fsDISCO (d), TDE 50% (e) and 70% (f), FAST9 for 3 (g) and 5 days (h), ClearSee for 3 (i) and 5 (j) days, ClearSee alpha for 3 (k), 5 (l), 10 (m) and 15 (n) days and CHAPS Clear for 3 (o), 5 (p), 10 (q) and 15 (r) days. Scale bar = 200 μm.

As anticipated, different clearing methods can remove tannins (PA) to a different extent. FAST9, CHAPS clear, and ClearSee alpha provided a reduced pink coloration (Fig. 1g, h, k-r). However, the clearing duration and efficiency of these methods were different. FAST9 removed PAs within 3-5d of incubation (Fig. 1g, h). For CHAPS clear and ClearSee alpha, it took about 15d to remove most of the PAs (Fig. 1k-r). However, we did observe pink staining around the embryo after clearing for 15d, which was still retained after 20d of clearing (not shown).

ClearSee, on the other hand, oxidized the PAs and caused formation of brown pigment (Fig. 1i, j). We did not prolong the incubation for more than 5d as the formation of brown pigment interfered with the imaging of Renaissance-stained seeds (Supplementary Fig. 2). The fsDISCO and 50% TDE methods did not prevent PA oxidation, as all seeds developed a pink coloration after vanillin staining (Fig. 1d, e). However, 70% TDE clearing resulted in a mix of pink (vanillin positive) and brown seeds (Fig. 1f). The presence of brown seeds could be due to oxidation. Similar results were observed in a negative control experiment with insufficient washing time (1h) of PFA-fixed seeds (Fig. 1c). Increasing the washing time improved staining uniformity in those samples, indicating that some chemicals may interfere with vanillin staining. The fsDISCO solution contains antioxidants, but the fsDISCO clearing protocol failed to remove PAs of cleared seeds, probably due to the shorter clearing time (Fig. 1d).

The vanillin staining results indicate that the presence of antioxidants is necessary to remove PAs or, at the least, to prevent their browning. The importance of antioxidants is demonstrated in Supplementary Fig. 3. In the absence of antioxidants such as sodium sulfite, PAs are oxidized in ClearSee and FAST9- (a FAST9 solution without antioxidants) (Supplementary Fig. 3a, c). However, the presence of antioxidants prevented PA oxidation and seed browning in ClearSee alpha and FAST9 with the addition of antioxidants (Supplementary Fig. 3b, d).

### Spectral properties of seeds treated with the different clearing methods

Since the vanillin staining indicated that the tested methods provide different removal levels and oxidation of the PAs, we assumed those can be associated with different spectral properties. Spectral characteristics of the cleared seeds compared to those of fluorescent proteins would then facilitate the selection of suitable clearing method.

We have used four excitation wavelength channels (405, 488, 514, and 561 nm) for the emission spectral profile of cleared seeds. Those channels are commonly used in LSCM (laser scanning confocal microscopy) imaging. Fresh seeds in PBS and PFA-fixed seeds were used as negative control (Fig. 2a, b). Both had a background level of autofluorescence in three (405, 488, and 561 nm) channels. They emitted fluorescence in the red emission range. fsDISCO displayed fluorescence in outer and inner integuments and embryo with 405 nm and 488 nm channels, whereas only in the inner integument and embryo with 561 nm excitation (Fig. 2c). In some seeds, we observed autofluorescence from endosperm nuclei (Supplementary Fig. 4). Seeds cleared with TDE predominantly emitted fluorescence in the red emission range, similarly to fixed seeds (Fig. 2d, e). TDE 70% showed higher autofluorescence, probably due to the seed collapse, increased refractive index, or both factors (Fig. 2e). ClearSee clearing induced some seed autofluorescence in the integuments in all excitation channels, with a strong autofluorescence in the red emission range after 561 nm excitation (Fig. 2f). This noticeable autofluorescence in the ii1 may be caused by the oxidized tannins. Seeds cleared with ClearSee alpha displayed some autofluorescence in the red emission range when excited at 405 and 561 nm (Fig. 2g), possibly due to oxidized tannins in ii1 that were not yet cleared within five days or other pigments present throughout the seed (Fig. 1l). Seeds cleared with FAST9 and CHAPS Clear displayed relatively low autofluorescence, especially with EasyIndex (EI) and 60% iohexol as mounting media (Fig. 2h-j). Of the four laser channels, the least seed autofluorescence was observed in the 514 nm excitation channel, indicating that using a YFP-based fluorescent reporter could be the best choice, irrespective of the clearing method used when considering the spectral properties of the cleared seeds.

**Fig. 2.**
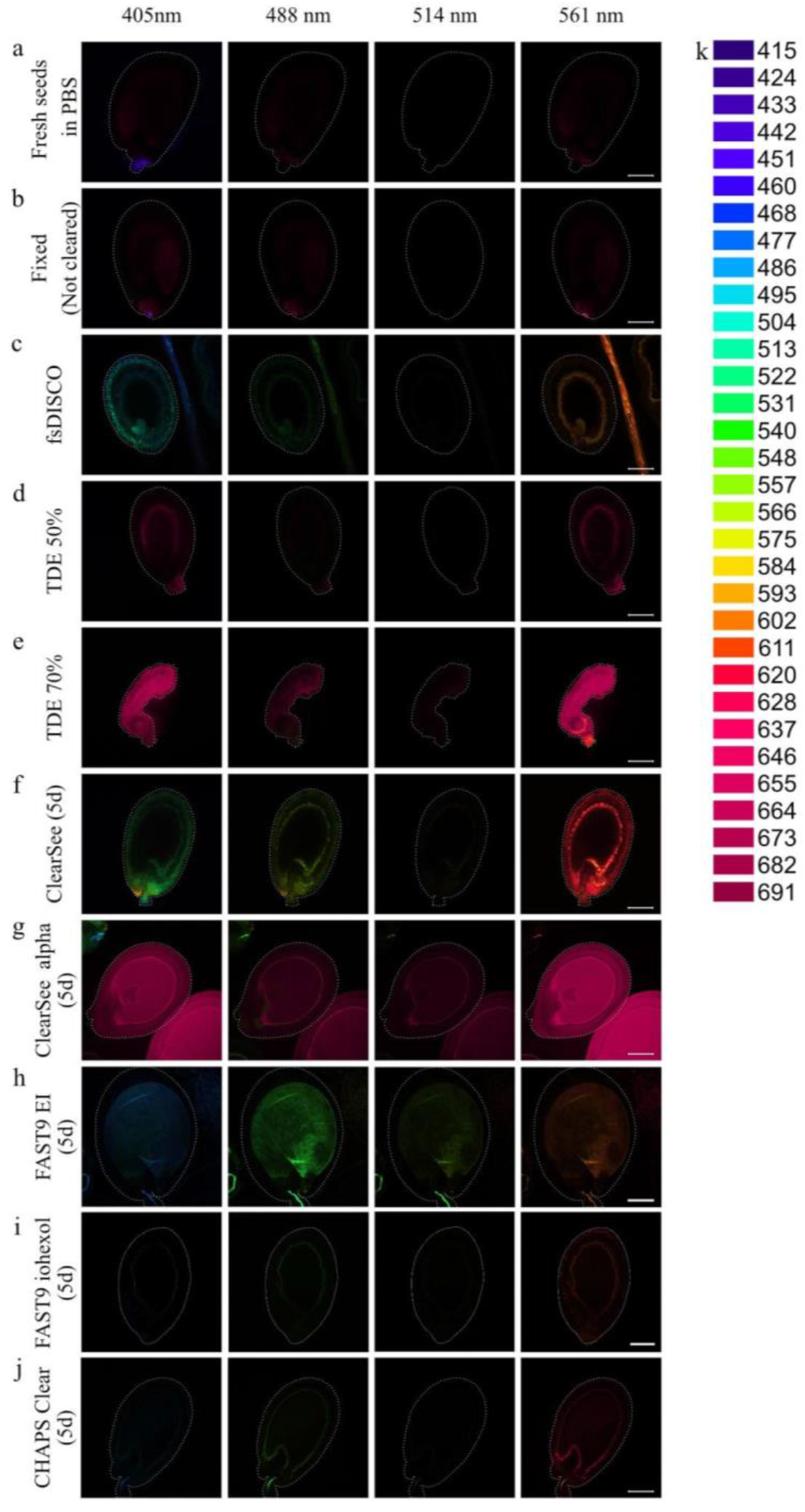
Spectral imaging of Arabidopsis seeds cleared with different clearing methods. The spectral imaging analysis was performed on fresh seeds in PBS (a), PFA-fixed not cleared seeds (b), seeds cleared with fsDISCO (c), TDE 50 % (d), TDE 70% (e), ClearSee (f), ClearSee alpha (g), FAST9 mounted in Easy Index (EI) (h), FAST9 mounted in iohexol (i) and CHAPS Clear (j). The seeds were cleared with ClearSee, ClearSee alpha, FAST9, and CHAPS Clear for five days. The emission spectra of the seeds for specific excitation channels (405, 488, 514, and 561 nm) were gathered. The collected emission spectra and their corresponding color code are shown (k). The seed perimeter is highlighted by a dashed line when the seed is not visible. Scale bar = 100 μm.

### The clearing protocols induced seed morphology changes

ClearSee and ClearSee alpha fluorescence-compatible clearing methods were used in ovules and seeds in the early development stages (Tofanelli et al. 2019; Imoto et al. 2021; Kurihara et al. 2021). Also, isolated embryos from the seeds up to torpedo stages were successfully cleared (Imoto et al. 2021). This study aimed at developing clearing methods for mature embryo-containing seeds. Our observations revealed that different methods differently impact seed morphology, depending on the embryonic developmental stage. In this experiment, we did not use the fsDISCO and TDE clearing methods, which resulted in high seed autofluorescence and seed collapsing, respectively (Fig. 2c-e).

Overall, ClearSee alpha and ClearSee had the least impact on the seed morphology, compared to the other tested methods (Fig. 3). The seeds with heart embryos, cleared with ClearSee alpha or ClearSee, frequently displayed undulating outer integuments (Fig. 3a, b, arrowheads).

**Fig. 3.**
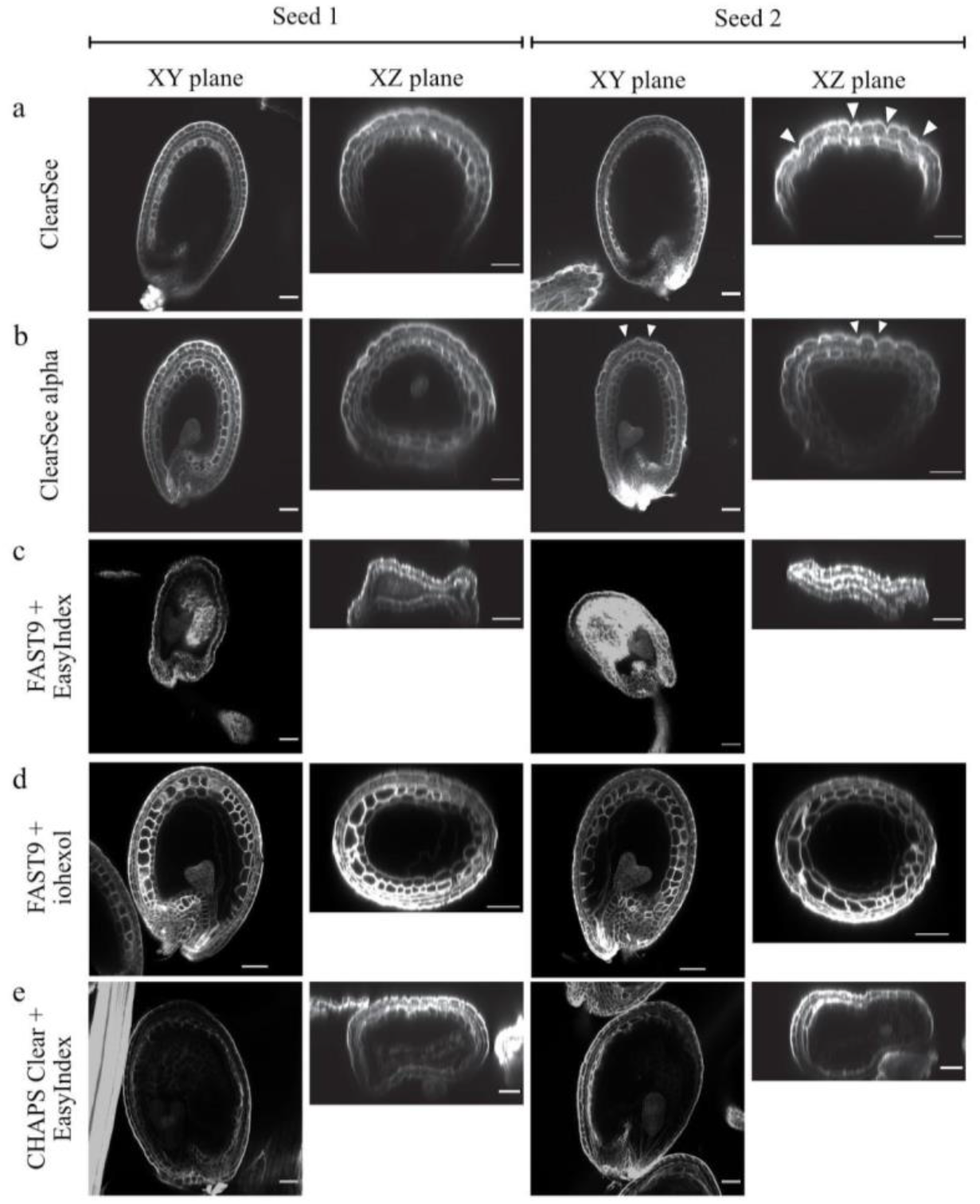
Comparison of morphological changes in heart-embryo-containing seeds induced by different clearing methods. ClearSee (a), ClearSee alpha (b), FAST9 mounted in 60% iohexol mounting medium, and CHAPS Clear with EasyIndex mounting medium (e) preserve seed morphology better than FAST9 combined with EasyIndex mounting medium (c). The imaging of two seeds per method is shown in the XY and XZ imaging axis. The arrowheads point to the undulation patterns in the outer integuments (a, b). The analysis of two representative seeds is presented. Scale Bar = 50 μm.

The most dramatic changes were observed in FAST9 and CHAPS-cleared seeds mounted in EI (Fig. 3c, e). The FAST9 + EI combination resulted in more severe seed collapses than the CHAPS clear + EI combination. To understand the contribution of EI to such morphological changes, we have used 50% TDE and PBS-T (Supplementary Fig. 5) as mounting media. In 50% of TDE-mounted seeds, the collapsing was not observed. However, irregular cell shape was observed in the integument cell layers (arrowheads in Supplementary Fig. 5c). PBS-T mounted seeds did not display any signs of deformations. Nevertheless, the clearing was not optimal in both cases (FAST9 + 50% TDE and FAST9 + PBS-T). The Renaissance staining signal was not detected in all layers of the inner integuments when comparing the seeds cleared by ClearSee and ClearSee alpha in Fig. 3a, b with those cleared with FAST9 (Fig. 2d, Supplementary Fig. 5). Recently, Sakamoto et al. (2022) used iohexol, a mounting medium with a high refractive index, to improve tissue transparency. We combined FAST9 clearing with 60% iohexol as a mounting medium to test whether it would better preserve seed morphology (Fig. 3d). Indeed, seeds were well cleared at the heart stage, and the seed morphology was preserved as well as with ClearSee alpha.

We observed drastic morphologic changes in seeds bearing embryos at the late globular to torpedo stages with samples cleared with TDE 70% (data not shown), FAST9, and CHAPS Clear (Fig. 3) when mounted with EI. Therefore, the impact of the seed developmental stage on the level of deformation associated with FAST9 + iohexol and CHAPS Clear was investigated in four embryonic developmental stages: 4/8-cell, 16-cell/early globular, heart, and mature stages (Fig. 4). The integuments display signs of shrinkage with both CHAPS Clear and FAST9 on XZ images that monitor clearing depth and seed deformity. The shrinkage is more pronounced in CHAPS and FAST9-cleared, EI-mounted seeds from the earliest tested developmental stage. The EI mounting medium appears to be responsible for the seed deformities, which is consistent with FAST9-cleared seeds in PBS-T mounting medium showing no signs of alteration at the cell or seed level (Supplementary Fig. 5). We tested 60% iohexol mounting medium in combination with FAST9 in seeds from the four development stages (Fig. 4 a-d). FAST9-cleared seeds mounted in 60% iohexol showed batch-to-batch variation in the clearing performance. The seed shape was mainly maintained. However, in most seeds, the endosperm cuticle detached, especially in the seeds with embryos from globular to torpedo stages where the endosperm is starting to cellularize (Fig. 3d). Nevertheless, this combination (FAST9 with 60% iohexol) provided the best preservation of the seed shape compared to other tested fast clearing of the tissues, except for ClearSee alpha.

**Fig. 4.**
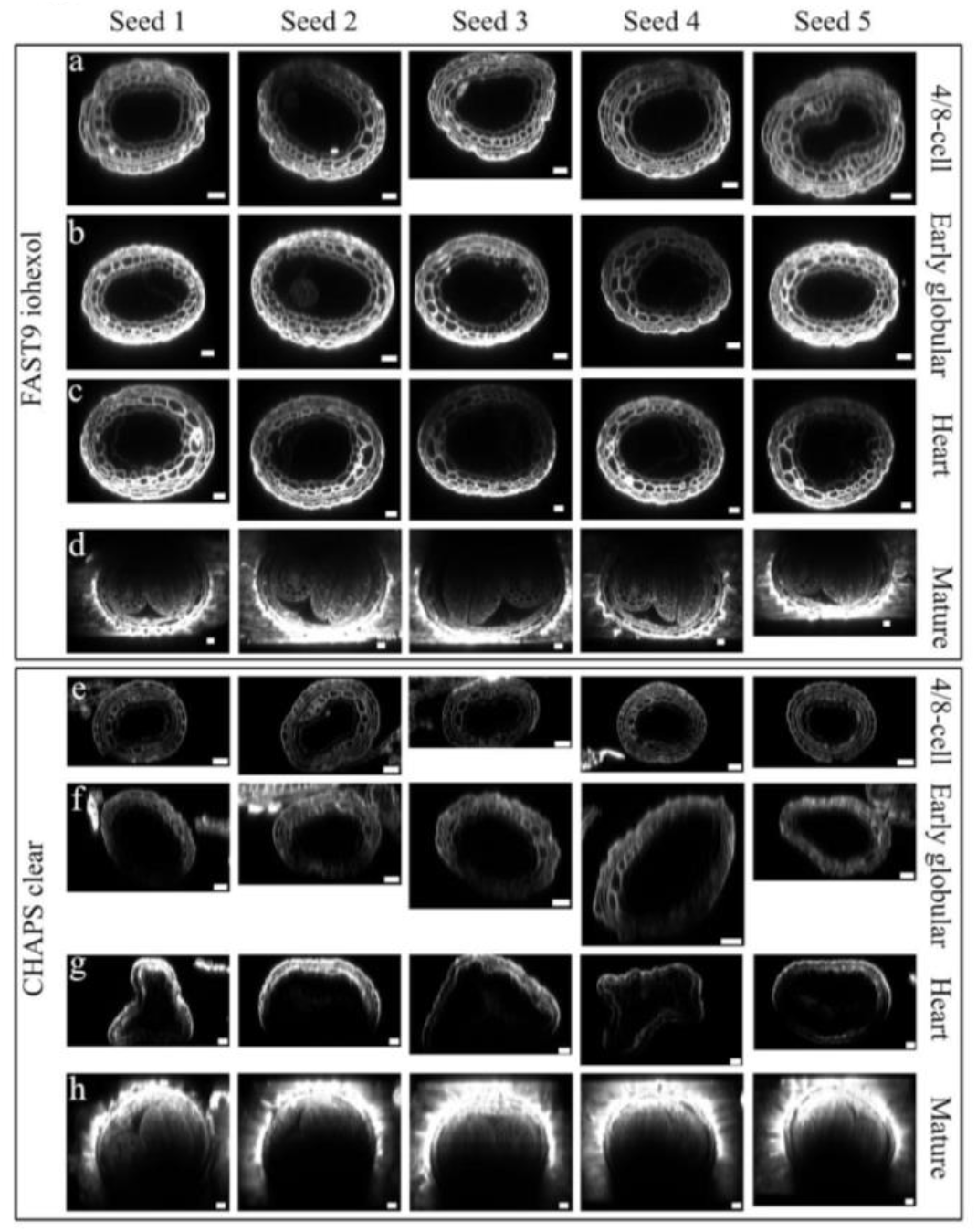
Impact of seed age on clearing-induced seed deformity. Arabidopsis seeds were cleared for five days with FAST9 and mounted in iohexol (a-d), or CHAPS Clear and mounted in EasyIndex (e-h). Spacers were used not to deform the seeds during mounting. Seeds with embryos at 4-8 cell (a, e), 16-cell/early globular (b, f), heart (c, g), and mature (d, h) stages were cleared and imaged. Five seeds are shown per method and growth stage as XZ images to show the impact on seed morphology. Scale Bar =25μm.

### ClearSee alpha preserves the fluorescence of FPs in cleared seeds up to the mature embryonic developmental stage

We established that FAST9 and CHAPS Clear clearing methods removed tannins (Fig. 1) and limited seed autofluorescence (Fig. 2), but did not preserve seed shape during the clearing protocol when associated with EI as a mounting medium (Figs 3 and 4). FAST9 associated with iohexol combines tannin removal, limited seed autofluorescence and a better seed shape preservation (Figs 1-4). We also demonstrated that while ClearSee was unable to remove tannins, resulting in a strong seed autofluorescence (Figs 1 and 2), ClearSee alpha removed them after a long clearing treatment (Fig. 1), which rendered the seeds non-autofluorescent when using the 488 and 514 nm excitation channels (Fig. 2). Also, ClearSee alpha had a limited impact on the seed shape (Fig. 3). Therefore, we applied the ClearSee alpha and FAST9 with iohexol clearing protocols on seeds from plants expressing the *pRPS5A::H2B-sGFP, pDR5::nVENUS* and *pWOX5::H2B-2xCHERRY* fluorescent reporters (Fig. 5).

**Fig. 5.**
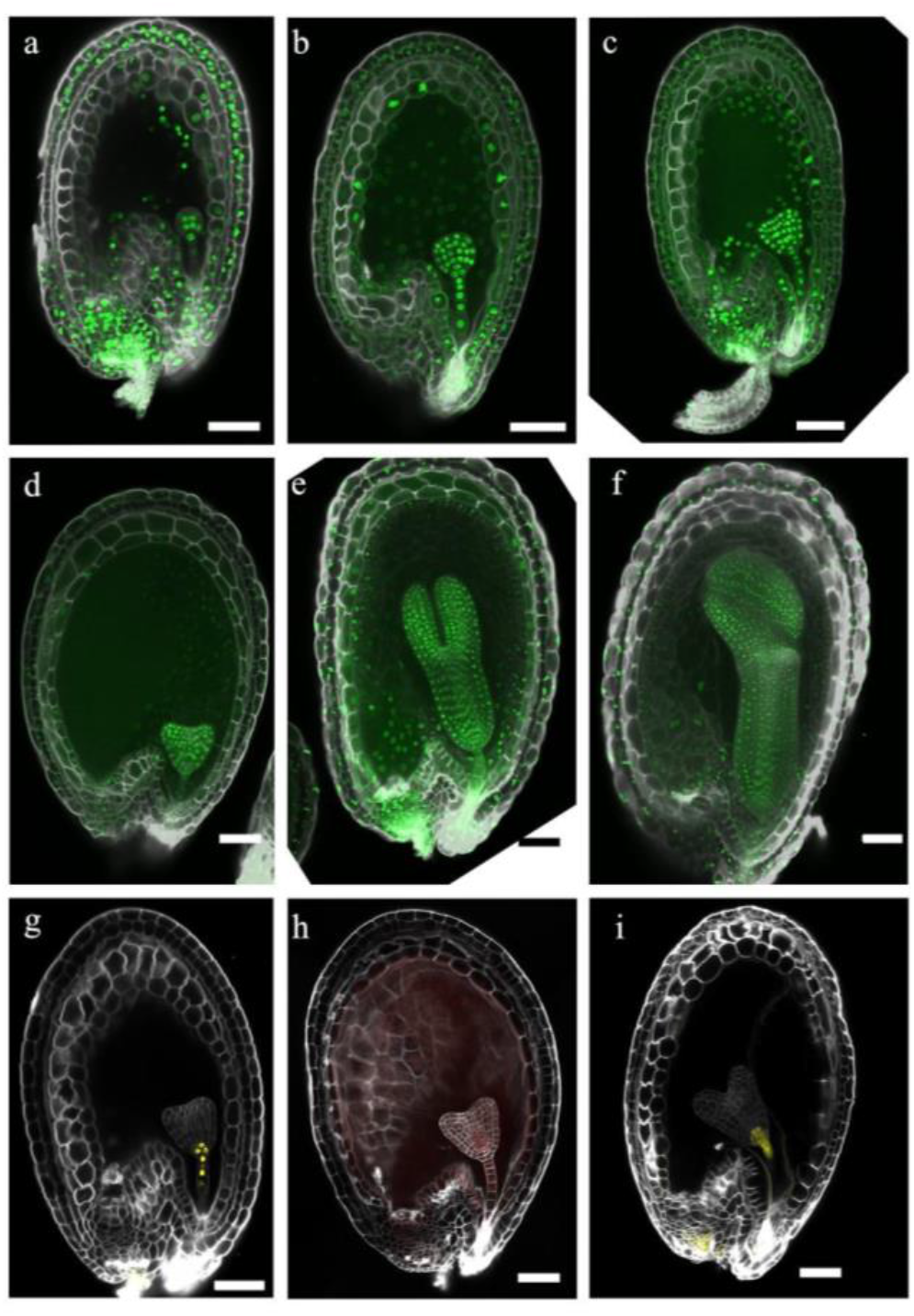
ClearSee alpha and FAST9 with iohexol preserve seed shape and fluorescence in mature seeds. (a-f) Seeds expressing *pRPS5A::H2B-sGFP* at early (a) and late globular (b), transition (c), heart (d), cotyledon (e) and torpedo (f) embryonic stages. The H2B-sGFP signal is in the nucleus as green fluorescence. (g, h) Seeds expression *pDR5::nVENUS* (g, i) and *pWOX5::H2B-2xCHERRY* (h) at heart embryonic stages. The yellow fluorescent VENUS and the CHERRY signals are targeted to the nucleus. The cell walls are counter-stained with Renaissance. Seeds were cleared with ClearSee alpha (a-h) or FAST9 + 60% iohexol (i). Scale bars = 50 μm.

ClearSee alpha cleared *pRPS5A::H2B-sGFP* seeds from early globular to torpedo embryonic stages while preserving the seed shape and GFP fluorescence in the embryo, endosperm, and integuments (Fig. 5a-f). Similarly, cleared seeds expressing *pDR5::nVENUS* and *pWOX5::H2B-2xCHERRY* were imaged from the heart to the mature stages (Fig. 5g, h for heart stage). As previously reported, VENUS fluorescence was observed in the embryo and the integuments (Robert et al. 2013; Figueiredo et al. 2016). The red fluorescence from the CHERRY protein was observed strongly in the quiescent center cells (as reported in (Haecker et al. 2004)) and in the inner and suspensor cells in heart-staged embryos. However, the mucilage in mature seeds interfered with seed clearing creating a fluorescent halo (not shown). We tested FAST9 clearing with mounting in EI on *pRPS5A::H2B-sGFP* seeds (Supplementary Fig. 6). We observed batch-to-batch inconsistency in the clearing performance and seed shape preservation. Most preserved seeds would have damaged or lost their outer integument cells during clearing. Gentle incubation at slightly elevated temperatures limited the seed collapse to occasional detachment of the endosperm cuticle and its collapse into the endosperm cavity. In addition, autofluorescence with the 488 nm excitation wavelength was observed in the vascular strands in the chalaza pole. The nuclear-targeted VENUS fluorescent signal of the *pDR5::nVENUS* seeds cleared with FAST9 and mounted in iohexol was slightly more diffuse when compared with *pDR5::nVENUS* seeds cleared with ClearSee alpha (Fig. 5i).

Similarly, we tested the CHAPS Clear clearing protocol on *pRPS5A::H2B-sGFP* and *pDR5::nVENUS* seeds (Supplementary Fig. 7). Seed shape deformities and shrinkage were observed. However, the clearing method mostly preserved the fluorescence of VENUS and GFP proteins. In mature seeds, the seed mucilage also interfered with the clearing, resulting in a fluorescent halo (Supplementary Fig. 7d).

This experiment indicated that ClearSee alpha remains the best clearing method that preserves the fluorescent signal of GFP, VENUS, and mCHERRY proteins in seeds bearing embryos from early globular to torpedo stages.

### ClearSee alpha-cleared seeds as a method to study embryo morphology

Seeds expressing *pDR5::nVENUS* and *pYUC1::3nGFP* were cleared using ClearSee alpha. Renaissance as a dye to counter-stain cell walls defines cell shape. After LSCM imaging, z-stack series were analyzed using Imaris 9.9.1 image processing software. Scans of seed coat structures were filtered out to focus on the embryo. Images were processed to generate a three-dimensional representation of the embryo (Fig. 6, Additional Movies 1 and 2). The starting material was a whole seed rather than an isolated embryo. Therefore, the embryonic structures were not damaged or flattened. Such analysis allows for an expression pattern analysis in three dimensions and an analysis of the cellular morphology of the embryo. It may be applied for detailed phenotyping and expression analysis of proteins of interest. For example, The *DR5* promoter is driving the expression of *VENUS* in the suspensor cells of the globular embryos and seed integuments (Fig. 6a-d). Also, the expression of the *YUC1* reporter is visible only in the protoderm cells (Fig. 6e,f).

**Fig. 6.**
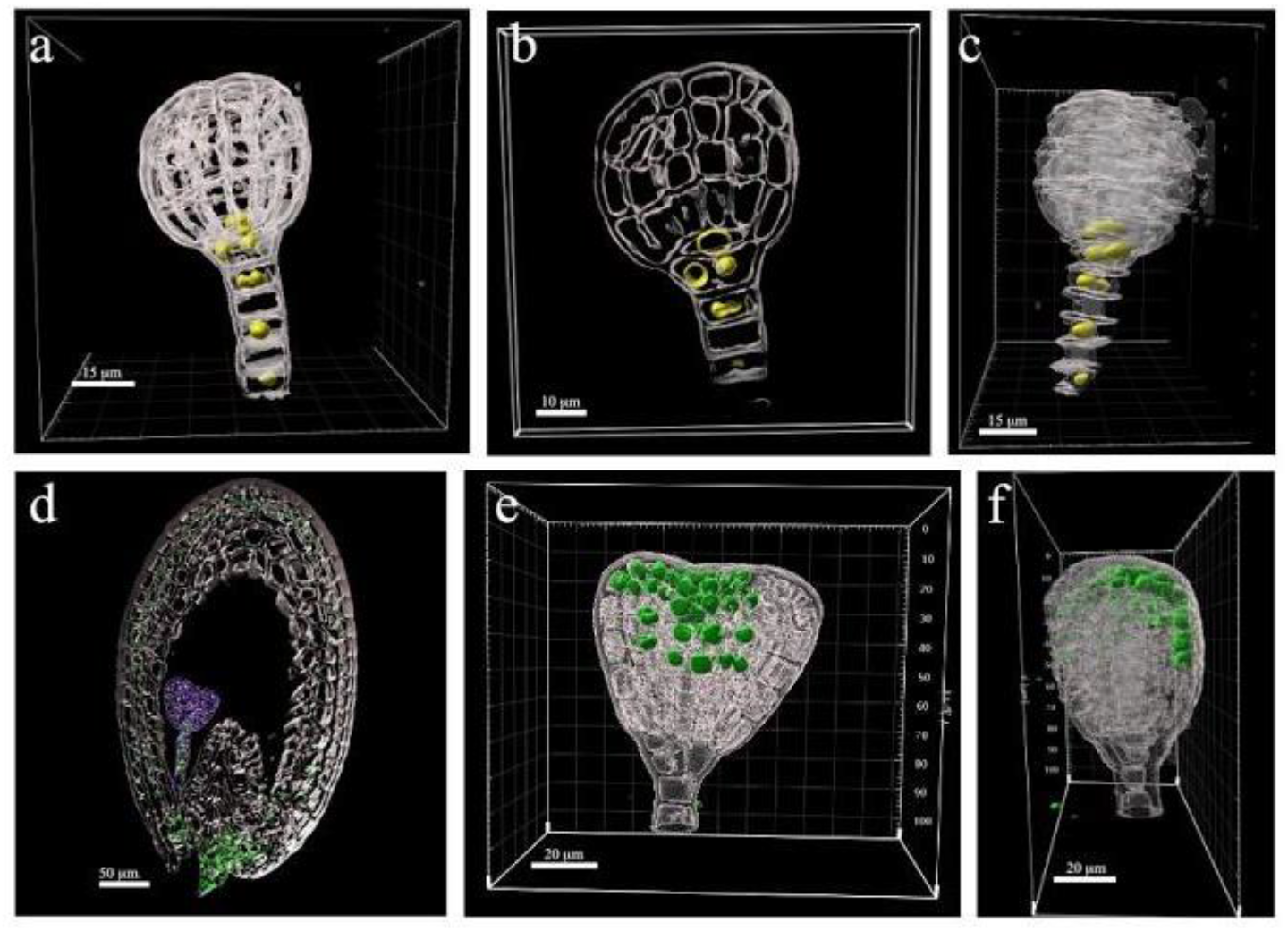
Three-dimensional representation of the embryos cleared within the seed. (a-c) Globular embryo expressing *pDR5::nVENUS*, in yellow: 3D view (a), middle section (b), side view (c) of one embryo, cropped out of the seed. (d) Whole seed expressing *pDR5::nVENUS*. in green. The embryo is marked in purple (e, f). Heart-shaped embryo expressing *pYUC1::3nGFP*, in green: 3D view (d) and side view (e) of an embryo cropped out of the seed. Images were processed with Imaris. The images of seed (d) and the embryo with *YUC1* pattern (e, f) were extracted from Additional Movies 1 and 2. The cells are marked in gray (Renaissance counter-stain). Scale bars as indicated in the panels.

## Discussion

An ideal clearing method allows for the clearing of tissues and organs in a short time, preserves their morphology, limits auto-fluorescence, and is compatible with the presence of dyes and fluorescent proteins. However, each existing method has its advantages and drawbacks, depending on the clearing mechanism and tissue being cleared. The clearing methods based on aqueous solutions provide slow clearance, while those utilizing organic solvents are more rapid and efficient. This study tested six methods, including one organic solvent-based clearing method, fsDISCO, on the Arabidopsis seeds of various developmental stages. The Arabidopsis seeds consist of tissues, pigments, storage material, and apoplastic barriers (Supplementary Fig. 1). The variation in the seed composition causes some methods to be more suitable than others.

fsDISCO is the only organic solvent-based clearing method we tested. It cleared the seed tissues within a day but increased their autofluorescence. There were also issues with the reproducibility of the method, and some seeds were morphologically preserved while others collapsed. In addition, this clearing method was incompatible with the Renaissance 2200 dye used to visualize cell walls. For these reasons, fsDISCO was not selected as the clearing method of choice.

TDE clearing was reported to rapidly clear seeds until globular stages (Musielak et al. 2016). However, in our study this method did not sufficiently clear the seeds beyond the globular stage and increasing the TDE percentage resulted in tissue disruption and seed collapse. Even if the tissue permeability of TDE is solved to preserve the tissue morphology, high TDE concentrations are not compatible with preservation of the fluorescence of some FPs (Musielak et al. 2016).

ClearSee and ClearSee alpha are related methods with similar protocols and outcomes, except for the presence of the sodium sulfite antioxidant in ClearSee alpha. The presence of antioxidants prevents the browning of PAs, resulting in a better optic penetration of the 405 nm emission laser. Oxidized tannins, as in ClearSee-cleared samples, interfere with imaging. First, oxidized tannins are known to absorb lower wavelengths, which cast shadows on the objects in the light path (Supplementary Fig. 2). Also, they produce autofluorescence interfering with the fluorescence signals produced by dyes or FPs that share the same emission/excitation laser channels (Fig. 2). In addition, ClearSee and ClearSee alpha clearing protocols take longer to clear tissue than fsDISCO and FAST9 (Table 1). However, these methods better preserve the seed morphology at all tested developmental stages.

**Table 1.** Comparison of the clearing methods in Arabidopsis seeds. DBE: Dibenzyl ether; EI: EasyIndex; RI: refractive index; d.n.s.: data not shown; n.a.: not available Clearing times are indicated, excluding the fixation duration. For ClearSee and CHAPS Clear, the clearing is better with more extended incubation periods.

| Protocol | Duration | Mounting medium | Mounting medium RI | tannin removal | autofluorescence | seed shape preservation | fluorescence preservation |
| --- | --- | --- | --- | --- | --- | --- | --- |
| fsDISCO (solvent-based) | 1d | DBE | 1.562 | -- | high, integuments | sporadic | moderately preserved (d.n.s.) |
| TDE 50% | 3-6h | TDE 50% | ~1.43 | -- | limited | mostly preserved (d.n.s.) | moderately preserved (d.n.s.) |
| TDE 70% | 3-6h | TDE 70% | ~1.47 | -- | strong | seed collapse | sporadic (d.n.s.) |
| ClearSee | 3-5d | ClearSee | 1.41 | -- (oxidation) | strong | mostly preserved | preserved |
| ClearSee alpha | up to 21d | ClearSee alpha | n. a. | ++ (15d) | limited | mostly preserved | preserved |
| FAST9 | 3-10d | EI | 1.52 | +++ | limited | seed collapse | preserved |
| FAST9 | 3-10d | TDE 50% | 1.43 | +++ | limited | mostly preserved | preserved |
| FAST9 | 3-10d | PBS-T | n. a. | +++ | limited | mostly preserved | preserved |
| FAST9 | 3-10d | 60% iohexol | 1.49 | +++ | limited | preserved | preserved |
| CHAPS Clear | 3-5d | EI | 1.52 | ++ (15d) | limited | seed collapse | preserved |

While FAST9 efficiently removes the seed tannins within three days, it does not preserve the seed morphology. We hypothesized that tissue preservation might depend on the mounting medium and tested FAST9 compatibility with the four mounting media. Combining FAST9 and EasyIndex worsened the morphological damage in the cleared sample. The incompatibility of EI with FAST9 may come from the slow EI diffusion into the seed tissues and the water removal from the cleared seeds. Mounting in 50 % TDE resulted in limited preservation of the seed morphology. PBS-T best preserved the seed morphology, but limited the laser penetration resulting in a reduced sharpness of images. Recently, iohexol was tested as a mounting medium in the TOMEI clearing method (Sakamoto et al. 2022). Iohexol has a high refractive index and appears to be a good mounting solution to combine with FAST9 to efficiently clear Arabidopsis seeds while mostly preserving the seed shape and fluorescence of FPs.

We noticed reproducibility and batch inconsistency issues with FAST9 clearing and EI or iohexol mounting, with some seeds with less to no collapse. In such cases, the outer layers of the seed coat were mechanically damaged during clearing (Supplementary Fig. 6). These observations suggest that the seed apoplastic barriers (epidermal and endosperm cuticles, Supplementary Fig. 1) could limit the diffusion of the EI mounting medium, causing the seed to collapse. Although the EI mounting medium is used in CHAPS Clear clearing, the morphological changes in CHAPS-cleared seeds were not as drastic as in the FAST9-cleared ones. Strong detergent (SDS) is used in FAST9 clearing, which could weaken the mechanical support of seed integument leading to the breaking of the tissue.

Irrespective of these drawbacks, FAST9 is suitable for clearing seeds, especially in case of older embryonic stages and morphology studies. This method could be especially helpful if the cuticles and other diffusion-limiting apoplastic barriers could be removed using enzymatic processes. In ClearSee and ClearSee alpha samples, the seed morphology is preserved mainly due to combining the clearing solution with the mounting medium having a low refractive index. Such a combination allows for a better diffusion of both solutions into the seed tissue before any mechanical damage to the integuments. Thus, ClearSee alpha appears to be the best suitable method for analysis of expression patterns using fluorescent proteins.

The tannin removal efficiency is very comparable in both CHAPS Clear and ClearSee alpha protocols, probably because both methods use similar detergents. However, CHAPS Clear loses its advantage upon its combination with the EI mounting medium, which affects the seed morphology. Finding a better solution than EI for mounting could therefore enhance the use of CHAPS Clear in seed clearing, notably as CHAPS Clear better preserves FPs’ fluorescence and has a high refractive index. Iohexol may be one of the mounting solutions that would solve this problem.

## Conclusion

In this study, we have compared six methods, of which three were new (fsDISCO, FAST9, and CHAPS Clear), and three were already reported (ClearSee, ClearSee alpha, TDE). Unlike previous studies limiting their analysis to pistils and seeds at early developmental stages, we tested those clearing methods in seeds from the 4/8-cell to the mature embryonic stage. We demonstrate that ClearSee alpha is the best clearing medium in such tissues as it preserves morphology and prevents oxidation of tannins. FAST9 is faster and performs better clearing of older seeds; however, it requires a suitable mounting medium. Seeds possess two apoplastic barriers and embryonic sheath, which complicate the diffusion of the clearing and mounting solutions. Removing such apoplastic barriers could improve seed clearing as seed tissues may become compatible with a greater variety of clearing methods suitable for deeper imaging.

## Methods

### Plant growth conditions

Arabidopsis seeds were sterilized using ethanol. After washing, seeds were sown in Murashige and Skoog (MS) medium plates containing 1% sucrose. Two days after cold stratification, plates were moved to a cultivation chamber (Photon Systems Instruments, CZ): 21/18°C (light/dark) with an 18-hour light/6-hour dark condition (LED illumination) and 50% humidity. After one week, seedlings were transferred to soil in individual pots. The plants were cultivated in phytotron chambers (Photon Systems Instruments, CZ) with the same conditions. In this study, Col-0, *pDR5::nVENUS* (Heisler et al. 2005; Wabnik et al. 2013), *pWOX5::H2B-2xCHERRY* (N2106156, SWELL Red Tide WOX5 line, (Marquès-Bueno et al. 2016) and *pRPS5a::H2B:sGFP* (a gift from Kurihara’s lab) were used.

### Fixation

The seed fixation was carried out using a fixative solution (4% Paraformaldehyde in 1X PBS with 0.05% Triton X100 [PBS-T]). Siliques at specific embryonic developmental stages were split open in 1X PBS-T and incubated in a 1.5 mL fixative in a microcentrifuge tube. The tubes were covered with aluminum foil throughout the procedure to be light-tight. The samples were vacuum infiltrated for ~30 min on ice to remove air bubbles. Then, the samples were incubated overnight at 4°C with slow rotation. The following day, samples were washed three times with 1X PBS-T at room temperature for one hour each.

### Vanillin staining

Vanillin staining is carried out as previously described (Xuan et al. 2014). Fixed and washed samples were dissected. Seeds were stained using freshly made 2 mL 1% vanillin solution (w/v, dissolved in 6N HCl) for one hour at room temperature with gentle rotation. Samples were mounted using the staining solution as a mounting medium. Visualization was performed immediately using a ZEISS Axioscope.A1 equipped with DIC optics, 10x objective, and an Axiocam 506 CCD camera, using bright field conditions. For cleared seeds, the staining was carried out after clearing and before the mounting steps.

### Renaissance SR2200 staining

A 0.1% (v/v) Renaissance SR2200 (Renaissance Chemicals, UK) solution was used to stain the samples. Since those clearing procedures involve organic solvents and to achieve uniform staining, samples were stained before clearing for fsDISCO and TDE clearing by adding the Renaissance dye into the washing solutions after fixation. On the contrary, seeds were stained post-clearing for ClearSee, ClearSee alpha, FAST9, and CHAPS. In such a case, the staining step was performed after clearing either with PBS-T (FAST9 and CHAPS) or by mixing the dye with the mounting medium (ClearSee and ClearSee alpha). After staining, the samples were washed for one hour in the same solution used for the staining procedure.

### fsDISCO clearing

fsDISCO has two steps: dehydration and refractive index matching (Qi et al. 2019). Fixed samples were dehydrated in THF (tetrahydrofuran) solutions in a concentration series of 40%, 60%, 80%, 90%, and three times 100% (v/v), with one-hour incubation each. THF solutions were diluted with distilled water, and the pH was adjusted to 9.0 with 1:10 Triethylamine (in water). Pure DBE (dibenzyl ether, pH not adjusted) is used as a mounting medium to clear the tissue after dehydration and to match the refractive index. Samples were incubated for three hours in DBE. All incubations are performed at 4°C with slow rotation. Because THF and DBE may contain peroxides, which quench fluorescence, the solutions were purified and stored according to Becker et al. (2012)

### TDE clearing

The TDE (2,2’-thiodiethanol) clearing protocol is adapted from Slane et al. (2017). TDE solutions were diluted in deionized water at 50%, 60%, and 70%. Fixed samples were transferred to a TDE solution at the desired dilution (as indicated in the text) for 10 min. The solution is refreshed, and samples are incubated for 1 to 3 hours at room temperature with slow rotation. A higher percentage of TDE resulted in seed collapse after the incubation. We found that 60% TDE is a good compromise between a good clearing with minimal or no seed collapse.

### FAST9 clearing and EasyIndex mounting

Fixed samples were incubated in a clearing solution (300 mM sodium dodecyl sulfate, 10 mM boric acid, and 100 mM sodium sulfite, pH adjusted to 9 using NaOH) at 37°C with gentle rotation for 3 to 10 days, depending on the sample size. Cleared samples were washed with PBS-T and stained with Renaissance overnight at 37°C with gentle rotation. Samples were stored in fresh 1X PBS or index-matched with EasyIndex (3-6 hr.) before microscopy.

### FAST9 clearing and iohexol mounting

Cleared samples with FAST9, as described above, were mounted in iohexol with modifications on the final iohexol concentration (Sakamoto et al. 2022). In brief, after overnight washing with PBS-T, the cleared samples were incubated at room temperature in a concentration series of 20%, 50%, and 60% (w/w) iohexol (TCI) in PBS for 10 minutes each to avoid osmotic stress and seed collapse. The samples were incubated in 60% iohexol for one hour with gentle shaking. Samples were freshly mounted in 60% iohexol. Iohexol solutions were stored at 4°C.

### ClearSee clearing

Fixed siliques were cleared with a ClearSee solution (10% xylitol (w/v), 15% sodium deoxycholate (w/v), 25% urea (w/v) in water) as previously described (Kurihara et al. 2015). Fixed and washed samples were incubated in the ClearSee solution at room temperature with slow rotation for three to 15 days, depending on the experiment. The ClearSee solution was refreshed every day. The samples were stained with Renaissance at the last incubation, washed, and mounted in ClearSee for visualization.

### ClearSee alpha clearing

The ClearSee alpha solution was prepared according to Kurihara et al. (2021), e.g., ClearSee solution supplemented with 50 mM sodium sulfite as an antioxidant. The protocol is similar to the one used for ClearSee clearing. The clearing was extended to 21 days with gentle shaking at room temperature for seeds carrying heart-stage embryos and older stages.

### CHAPS clear clearing

The procedure is similar to the FAST9 clearing method. The clearing solution comprises 10% CHAPS solution (in water), 10 mM boric acid, and 100mM sodium sulfite, pH adjusted to 9 using NaOH. After clearing, the samples were washed three times with 1X PBS for one hour each. The seeds were mounted with EasyIndex.

### Mounting of cleared samples

Sample Mounting methods depend on the thickness of the samples. The samples were mounted in the respective mounting medium (as mentioned for each clearing method). For whole seeds, we used one or two coverslips (depending on the thickness of the sample) as spacers attached to the slide with Fixogum glue (Marabu). For cleared siliques, we used Press-to-Seal Silicone Isolators Adhesive (cat no 70336-70, Electron Microscopy Sciences). Depending on the sample thickness, 1-2 spacers were used.

### Microscopy

Fluorescence and spectral imaging were performed using Zeiss LSM 700, 780, and 880 confocal microscopes. Expression pattern analysis was performed on ZEISS 700 and 880 with a correction collar 25x magnification objective at 0.6 zoom at 1024×1024 pixel resolution. Two laser lines were used: 405 nm for imaging Renaissance combined with 488 nm for GFP and VENUS or 561 nm for mCHERRY imaging. Spectral images were taken on Zeiss LSM 780 with a correction collar 25× magnification objective at 0.6 zoom. Scans were performed in a 2048*2048 pixel resolution. For each seed, four laser lines (405, 488, 514, 561 nm) were used to capture the spectral properties. The collection of emitted light varies for each laser line: 411-695nm, 491-695nm, 517-695nm, and 562-695nm, respectively, with 8nm intervals. The images were captured in 12 bits. The laser intensity and detector gain were kept constant for samples. For monitoring seed collapsing, seeds were scanned using XY and XZ scanning mode. XY scanning was performed at 1024×1024 pixel mode. XZ scanning was done with system-optimized values to get the highest possible resolution. Excitation laser intensity (405 nm) was adjusted for each seed to the level that a few saturated pixels appeared. Images were captured with Zeiss LSM 780 with a correction collar 25x magnification objective at 0.6 zoom (for XY). Emission spectra were collected from 410 to 500 nm.

### Image analysis

Images were processed with ZEISS ZEN Blue and Imaris v.9.9.1 software and analyzed using ImageJ (Fiji). Image panels were prepared in Affinity Publisher.

## Acknowledgments

We acknowledge the CELLIM Core Facility supported by the Czech-BioImaging large RI project (LM2018129 funded by MEYS CR) and the Plant Sciences Core Facility of CEITEC Masaryk University for their support in obtaining the scientific data presented in this paper.

The authors thank Kurihara’s lab for providing the Arabidopsis seeds used in this study.

## Statements and Declarations

### Availability of data and materials

Data sharing does not apply to this article as no datasets were generated or analyzed in the study.

### Author Contribution Statement

VPSA and HSR conceived and designed the project. VPSA and JFSL conducted the experiments. VPSA, JFSL, and HSR analyzed the data. LM and KP contributed to the preparation of some chemicals. VPSA, JFSL, and HSR wrote the manuscript. All authors read and approved the manuscript.

## List of Abbreviations

CHAPS: 3-[(3-cholamidopropyl)dimethylammonio]-1-propane sulfonate.
DBE: dibenzyl ether
3DISCO: 3-dimensional imaging of solvent-cleared organs
EI: EasyIndex
FAST9: Free of acrylamide sodium dodecyl sulfate-based tissue clearing at pH9
FPs: Fluorescent proteins
GFP: green fluorescent protein
ii: inner integuments
LSCM: laser scanning confocal microscopy
PA: proanthocyanidins
PBS-T: phosphate buffer saline with tween
PFA: paraformaldehyde
SDS: sodium dodecyl sulfate
TDE: 2,2-thiodiethanol
THF: tetrahydrofuran
YFP: yellow fluorescent protein

## Supplementary Information

**Supplementary Fig. 1** An illustration showing the structure of heart-stage Arabidopsis seed with secondary metabolites (flavonol, tannins) and apoplastic barriers (epidermal and endosperm cuticles, embryo sheath, and suberin). The illustration is adapted from Verma, S et al. 2022.

**Supplementary Fig. 2** Arabidopsis seeds cleared (ClearSee and TDE) and stained with Renaissance SR2200

(a, b) Seeds cleared with ClearSee, (c, d) or TDE clearing methods. The presence of oxidized tannins is casting shadows on the inner walls of inner integument 1 (white arrows) and in the endosperm cavity (arrowheads in b). Image (d) was taken with a higher magnification to avoid saturation of the detector. Scale bar = 25μm.

**Supplementary Fig. 3** Antioxidants are necessary to prevent the browning of tannins Five-dap Arabidopsis seeds were cleared with clearing media without (a, c) or with (b, c) antioxidant. The seeds cleared without antioxidants have browning of the tannins. The clearing media used are (a) ClearSee, (b) ClearSee alpha, (c) FAST9 without antioxidants, and (d) FAST9 cleared with antioxidants. Scale bar = 200μm

**Supplementary Fig. 4** fsDISCO-cleared seeds displayed auto-fluorescence in cell walls and endosperm nuclei

The cleared seed was excited with 405 (a), 488 (b), 514 (c), and 561 (d) nm excitation channel and the emission spectra collected as mentioned in the methods. The color-coding of the emission colors is indicated in Figure 2j. Scale bars = 50μm.

**Supplementary Fig. 5** Arabidopsis seeds (heart stage) cleared with FAST9 clearing agent for three days and mounted in PBS-T (a, b) and TDE 50% (c, d)

A representative sample of two seeds is presented. No signs of cell shrinkage is visible as shown in the insets (a, b) or some signs of undulations in the inner integuments the XZ pictures (arrowheads, c, d). Seeds were cleared at room temperature and at 100 rpm to minimize the seed wall damage. Scale bar = 50μm.

**Supplementary Fig. 6** Arabidopsis seeds expressing *pRPS5a::H2B:sGFP* cleared for 5 days with FAST9 clearing agent and mounted in EasyIndex

The lack of seed collapsing was demonstrated with seeds of different developmental stages: (a) octant, (b) globular, (c) transition, (d-f) heart, and (g) torpedo stages. The arrowheads indicate the detachment of the endosperm cuticle from the integuments and its collapse into the endosperm cavity. Samples were cleared at 180 rpm at 37°C. Scale bar = 50μm.

**Supplementary Fig. 7** Arabidopsis seeds expressing *pDR5::nVENUS* (a-c) and *pRPS5a::H2B:sGFP* (d), cleared for 5 days with CHAPS Clear and mounted in EasyIndex. Scale bar = 50μm.

**Supplementary Movie 1**

Cleared seed with ClearSee alpha expressing *pDR5::nVENUS*.

Z-stacks were processed in Imaris.

Single section is visible in Fig. 6d.

**Supplementary Movie 2**

Cleared seed with ClearSee alpha expressing *pYUC1::3nGFP*.

Z-stacks were processed in Imaris.

The embryo was cropped out and isolated from the seed integuments for illustration purposes.

Single sections are visible in Figs 6e,f.

## References

Becker K, Jährling N, Saghafi S, et al (2012) Chemical Clearing and Dehydration of GFP Expressing Mouse Brains. Plos One 7:e33916. https://doi.org/10.1371/journal.pone.0033916

Debeaujon I, Nesi N, Perez P, et al (2003) Proanthocyanidin-accumulating cells in Arabidopsis testa: regulation of differentiation and role in seed development. The Plant cell 15:2514–2531. https://doi.org/10.1105/tpc.014043

Doll NM, Ingram GC (2022) Embryo–Endosperm Interactions. Annu Rev Plant Biol 73:293–321. https://doi.org/10.1146/annurev-arplant-102820-091838

Ertürk A, Becker K, Jährling N, et al (2012) Three-dimensional imaging of solvent-cleared organs using 3DISCO. Nat Protoc 7:1983–1995. https://doi.org/10.1038/nprot.2012.119

Figueiredo DD, Batista RA, Roszak PJ, et al (2016) Auxin production in the endosperm drives seed coat development in Arabidopsis. eLife 5:e20542. https://doi.org/10.7554/elife.20542

Haecker A, Groß-Hardt R, Geiges B, et al (2004) Expression dynamics of WOX genes mark cell fate decisions during early embryonic patterning in Arabidopsis thaliana. Development (Cambridge, England) 131:657–668. https://doi.org/10.1242/dev.00963

Hahn C, Becker K, Saghafi S, et al (2019) High-resolution imaging of fluorescent whole mouse brains using stabilised organic media (sDISCO). J Biophotonics 12:e201800368. https://doi.org/10.1002/jbio.201800368

Heisler MG, Ohno C, Das P, et al (2005) Patterns of auxin transport and gene expression during primordium development revealed by live imaging of the Arabidopsis inflorescence meristem. Current biology 15:1899–1911. https://doi.org/10.1016/j.cub.2005.09.052

Hériché M, Arnould C, Wipf D, Courty P-E (2022) Imaging plant tissues: advances and promising clearing practices. Trends Plant Sci 27:601–615. https://doi.org/10.1016/j.tplants.2021.12.006

Hoyer (1882) Beitrage Zur Histologischen Technik. Biologisches Zentralblatt Bd. 2:

Imoto A, Yamada M, Sakamoto T, et al (2021) A ClearSee-Based Clearing Protocol for 3D Visualization of Arabidopsis thaliana Embryos. Plants 10:190. https://doi.org/10.3390/plants10020190

Kurihara D, Mizuta Y, Nagahara S, Higashiyama T (2021) ClearSeeAlpha: Advanced Optical Clearing for Whole-Plant Imaging. Plant Cell Physiol 62:1302–1310. https://doi.org/10.1093/pcp/pcab033

Kurihara D, Mizuta Y, Sato Y, Higashiyama T (2015) ClearSee: a rapid optical clearing reagent for whole-plant fluorescence imaging. Development (Cambridge, England) 142:4168–4179. https://doi.org/10.1242/dev.127613

Marquès-Bueno MDM, Morao AK, Cayrel A, et al (2016) A versatile Multisite Gateway-compatible promoter and transgenic line collection for cell type-specific functional genomics in Arabidopsis. The Plant Journal 85:320–333. https://doi.org/10.1111/tpj.13099

Matilla AJ (2019) Seed coat formation: its evolution and regulation. Seed Sci Res 29:215–226. https://doi.org/10.1017/s0960258519000254

Musielak TJ, Slane D, Liebig C, Bayer M (2016) A Versatile Optical Clearing Protocol for Deep Tissue Imaging of Fluorescent Proteins in Arabidopsis thaliana. PLoS ONE 11:e0161107–e0161107. https://doi.org/10.1371/journal.pone.0161107

Palmer WM, Martin AP, Flynn JR, et al (2015) PEA-CLARITY: 3D molecular imaging of whole plant organs. Sci Rep-uk 5:13492. https://doi.org/10.1038/srep13492

Park Y-G, Sohn CH, Chen R, et al (2019) Protection of tissue physicochemical properties using polyfunctional crosslinkers. Nat Biotechnol 37:73–83. https://doi.org/10.1038/nbt.4281

Qi Y, Yu T, Xu J, et al (2019) FDISCO: Advanced solvent-based clearing method for imaging whole organs. Sci Adv 5:eaau8355. https://doi.org/10.1126/sciadv.aau8355

Robert HS, Grones P, Stepanova AN, et al (2013) Local auxin sources orient the apical-basal axis in Arabidopsis embryos. Current biology 23:2506–2512. https://doi.org/10.1016/j.cub.2013.09.039

Sakamoto Y, Ishimoto A, Sakai Y, et al (2022) Improved clearing method contributes to deep imaging of plant organs. Commun Biology 5:12. https://doi.org/10.1038/s42003-021-02955-9

Slane D, Bürgel P, Bayer M (2017) Plant Germline Development, Methods and Protocols. Methods Mol Biology 1669:87–94. https://doi.org/10.1007/978-1-4939-7286-9_8

Tofanelli R, Vijayan A, Scholz S, Schneitz K (2019) Protocol for rapid clearing and staining of fixed Arabidopsis ovules for improved imaging by confocal laser scanning microscopy. Plant methods 1–13. https://doi.org/10.1186/s13007-019-0505-x

Verma S, Attuluri VPS, Robert HS (2022) Transcriptional control of Arabidopsis seed development. Planta 255:90. https://doi.org/10.1007/s00425-022-03870-x

Wabnik K, Robert HS, Smith RS, Friml J (2013) Modeling framework for the establishment of the apical-basal embryonic axis in plants. Current biology 23:2513–2518. https://doi.org/10.1016/j.cub.2013.10.038

Warner CA, Biedrzycki ML, Jacobs SS, et al (2014) An Optical Clearing Technique for Plant Tissues Allowing Deep Imaging and Compatible with Fluorescence Microscopy. Plant Physiol 166:1684–1687. https://doi.org/10.1104/pp.114.244673

Xuan L, Wang Z, Jiang L (2014) Vanillin Assay of Arabidopsis Seeds for Proanthocyanidins. Bio-protocol 4:e1309. https://doi.org/10.21769/bioprotoc.1309

Zhao S, Todorov MI, Cai R, et al (2020) Cellular and Molecular Probing of Intact Human Organs. Cell 180:796–812.e19. https://doi.org/10.1016/j.cell.2020.01.030

