## Supplementary Information for "Comparing the efficiency of six clearing methods in developing seeds of *Arabidopsis thaliana*"

### Supplementary Figure 1

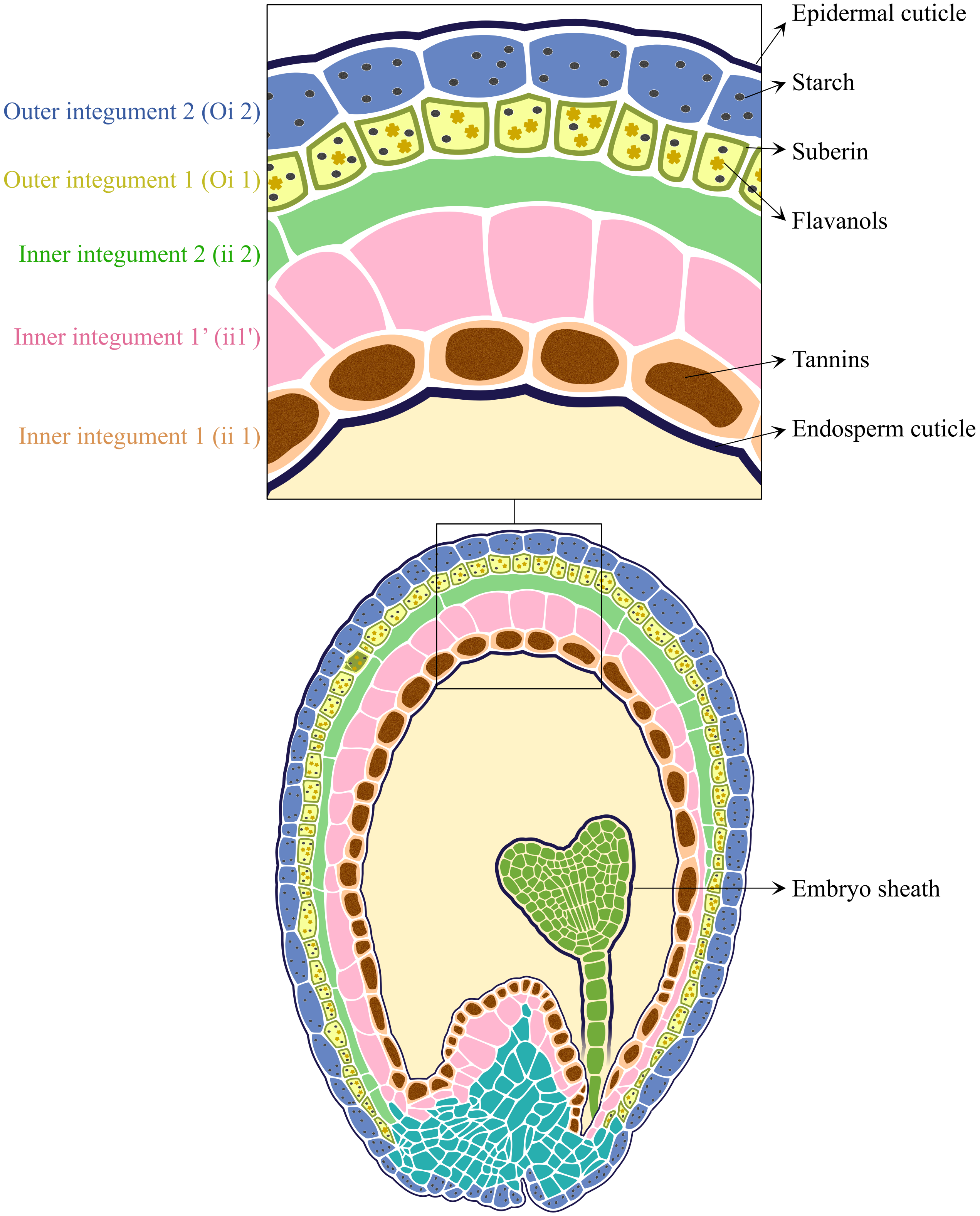


**Arabidopsis seed at Heart stage**

**Supplementary Fig. 1** An illustration showing the structure of heart-stage Arabidopsis seed with secondary metabolites (flavonol, tannins) and apoplastic barriers (epidermal and endosperm cuticles, embryo sheath, and suberin). The illustration is adapted from Verma, S et al. 2022.

### Supplementary Figure 2

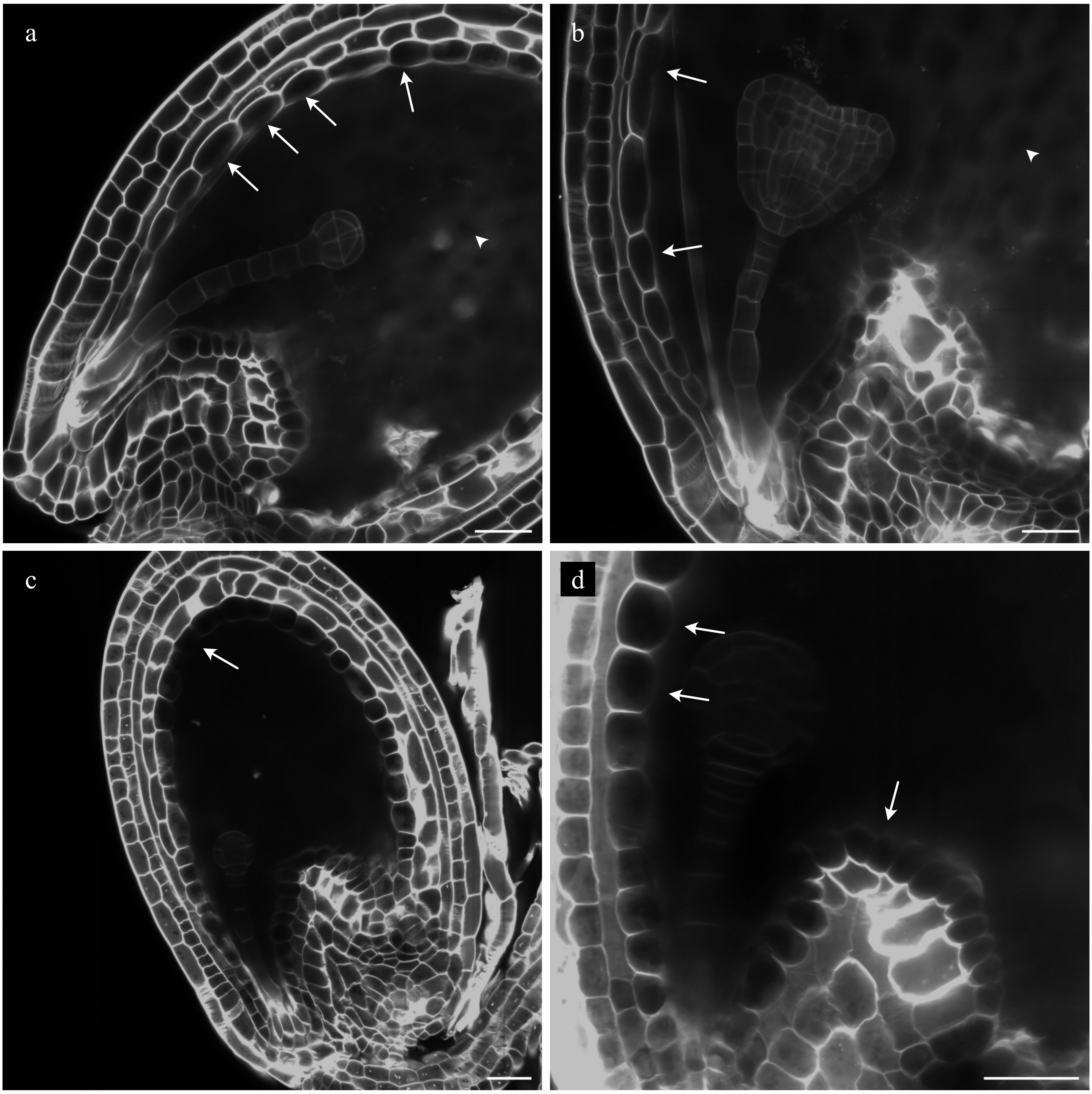


**Supplementary Fig. 2** Arabidopsis seeds cleared (ClearSee and TDE) and stained with Renaissance SR2200

(a, b) Seeds cleared with ClearSee, (c, d) or TDE clearing methods. The presence of oxidized tannins is casting shadows on the inner walls of inner integument 1 (white arrows) and in the endosperm cavity (arrowheads in b). Image (d) was taken with a higher magnification to avoid saturation of the detector. Scale bar = 25µm

### Supplementary Figure 3

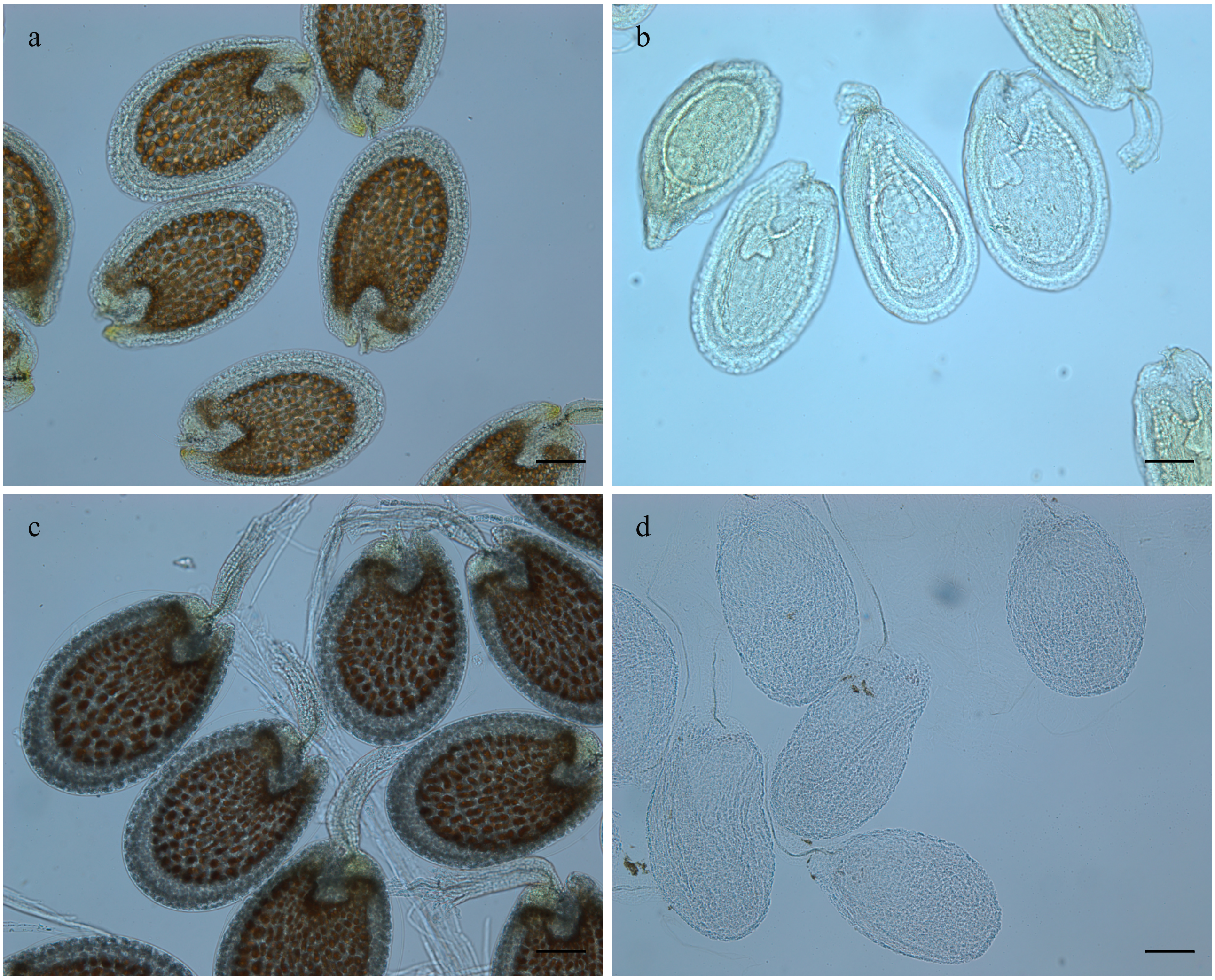


**Supplementary Fig. 3** Antioxidants are necessary to prevent the browning of tannins

Five-dap Arabidopsis seeds were cleared with clearing media without (a, c) or with (b, c) antioxidant. The seeds cleared without antioxidants have browning of the tannins. The clearing media used are (a) ClearSee, (b) ClearSee alpha, (c) FAST9 without antioxidants, and (d) FAST9 cleared with antioxidants. Scale bar = 200μm

### Supplementary Figure 4

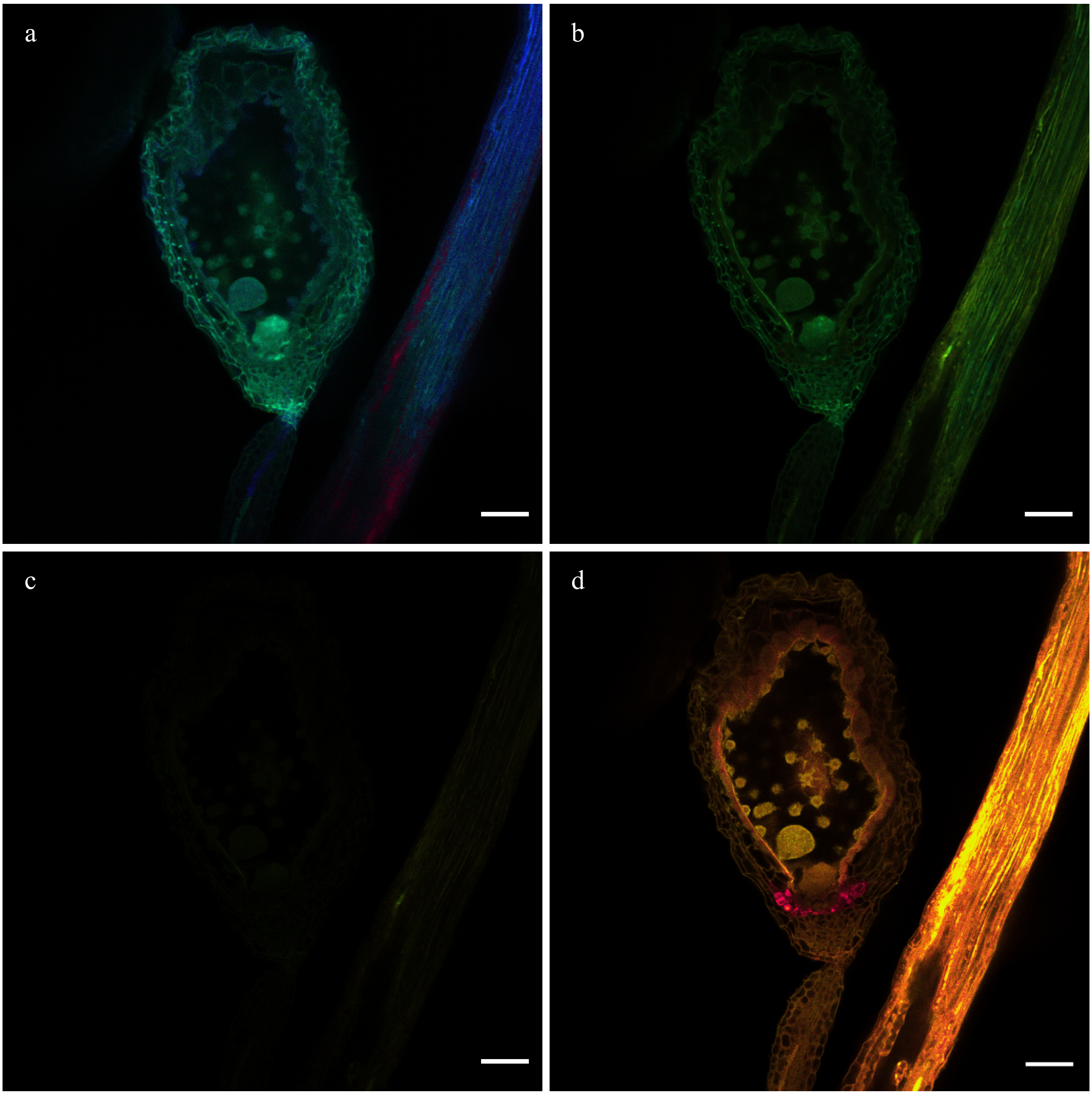


**Supplementary Fig. 4** fsDISCO-cleared seeds displayed auto-fluorescence in cell walls and endosperm nuclei

The cleared seed was excited with 405 (a), 488 (b), 514 (c), and 561 (d) nm excitation channel and the emission spectra collected as mentioned in the methods. The color-coding of the emission colors is indicated in Figure 2j. Scale bars = 50µm

### Supplementary Figure 5

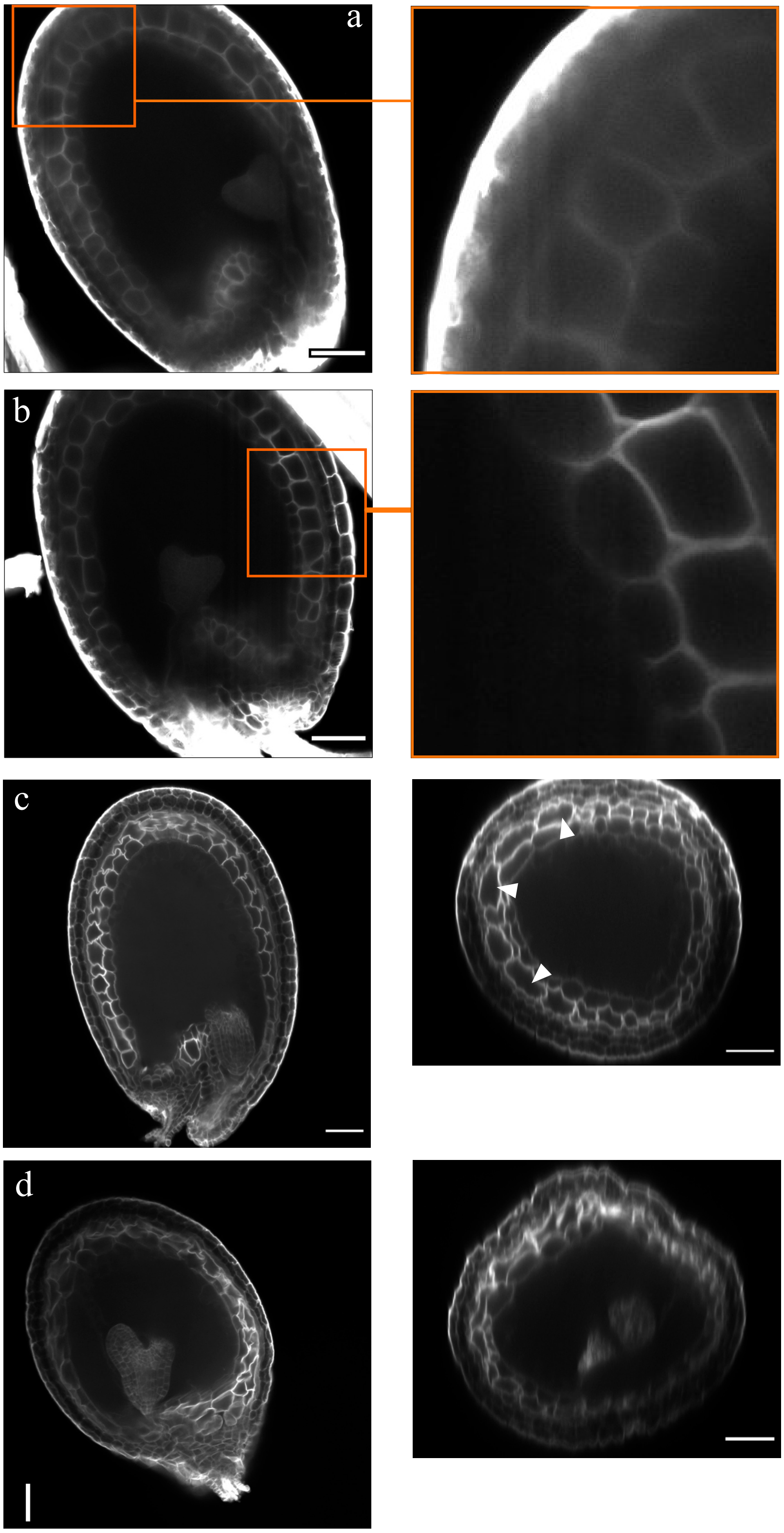


**Supplementary Fig. 5** Arabidopsis seeds (heart stage) cleared with FAST9 clearing agent for three days and mounted in PBS-T (a, b) and TDE 50% (c, d)

A representative sample of two seeds is presented. No signs of cell shrinkage is visble as shown in the insets (a, b) or some signs of undulations in the inner integuments the XZ pictures (arrowheads, c, d). Seeds were cleared at room temperature and at 100 rpm to minimize the seed wall damage. Scale bar = 50µm.

### Supplementary Figure 6

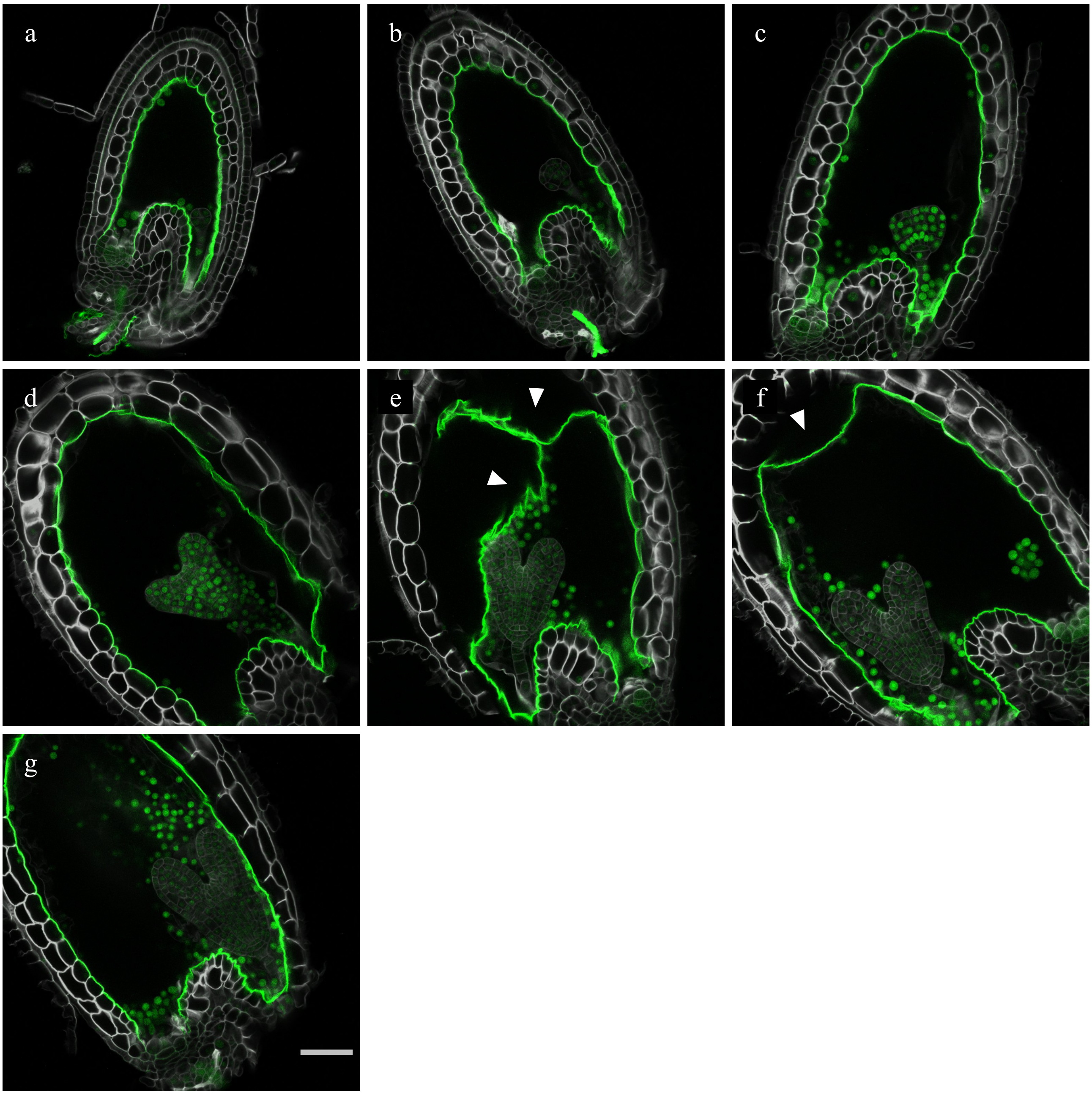


**Supplementary Fig. 6** Arabidopsis seeds expressing *pRPS5a::H2B:sGFP* cleared for 5 days with FAST9 clearing agent and mounted in EasyIndex

The lack of seed collapsing was demonstrated with seeds of different developmental stages: (a) octant, (b) globular, (c) transition, (d-f) heart, and (g) torpedo stages. The arrowheads indicate the detachment of the endosperm cuticle from the integuments and its collapse into the endosperm cavity. Samples were cleared at 180 rpm at 37°C. Scale bar = 50µm.

### Supplementary figure 7

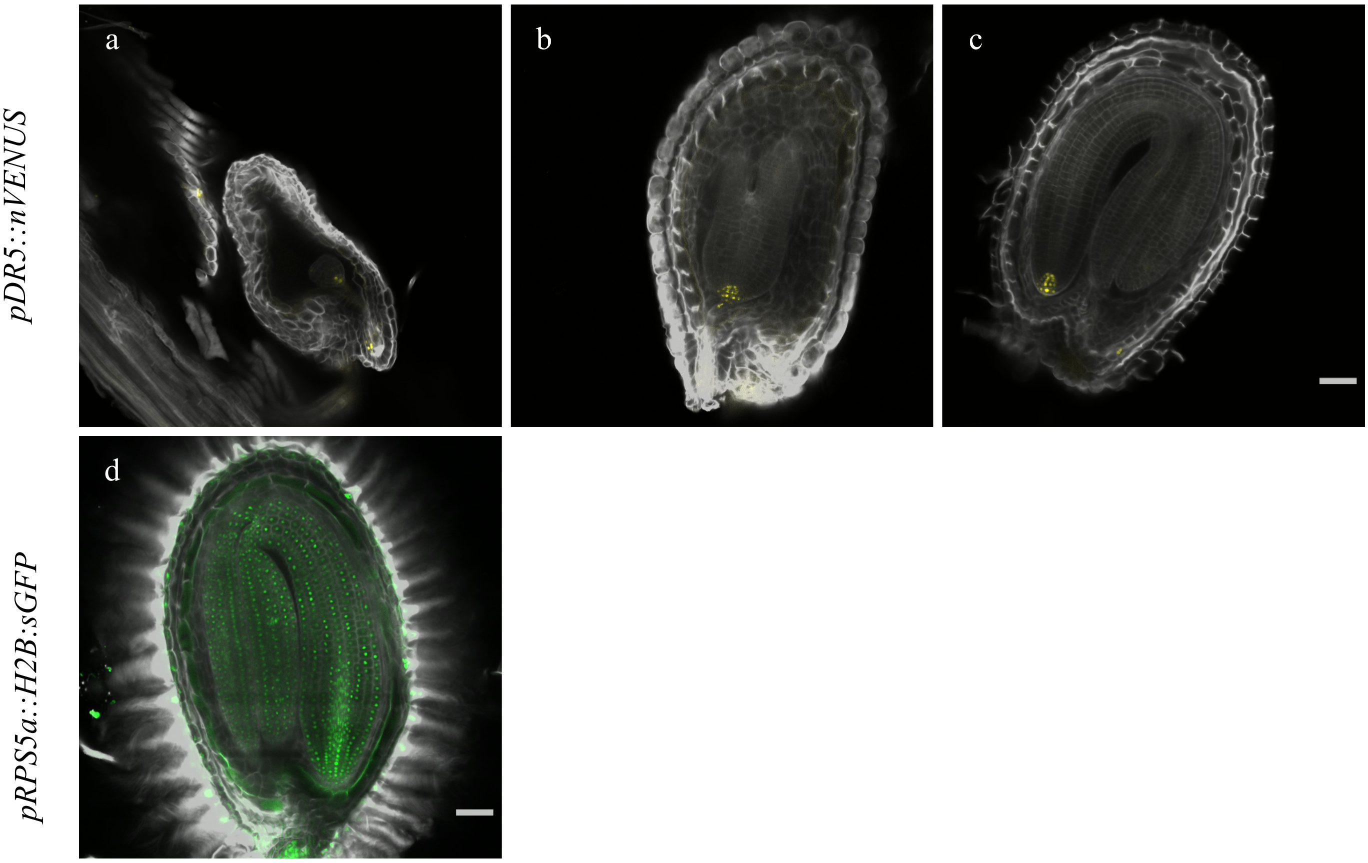


**Supplementary Fig. 7** Arabidopsis seeds expressing *pDR5::nVENUS* (a-c) and *pRPS5a::H2B:sGFP* (d), cleared for 5 days with CHAPS Clear and mounted in EasyIndex. Scale bar = 50µm.

**Supplementary Movie 1**
Cleared seed with ClearSee alpha expressing *pDR5::nVENUS*.
Z-stacks were processed in Imaris.
Single section is visible in Fig. 6d.

**Supplementary Movie 2**
Cleared seed with ClearSee alpha expressing *pYUC1::3nGFP*.
Z-stacks were processed in Imaris.
The embryo was cropped out and isolate from the seed integuments for illustration purposes.
Single sections are visible in Figs 6e, f.
